# Domains of methylated CAC and CG target MeCP2 to tune transcription in the brain

**DOI:** 10.1101/087577

**Authors:** Sabine Lagger, John C Connelly, Gabriele Schweikert, Shaun Webb, Jim Selfridge, Bernard H Ramsahoye, Miao Yu, Dina DeSousa, Christian Seiser, Chuan He, Guido Sanguinetti, Lawrence C Sowers, Malcolm D Walkinshaw, Adrian Bird

## Abstract

Mutations in the gene encoding the methyl-CG binding protein MeCP2 cause neurological disorders including Rett syndrome. The di-nucleotide methyl-CG (mCG) is the canonical MeCP2 DNA recognition sequence, but additional targets including non-methylated sequences have been reported. Here we use brain-specific depletion of DNA methyltransferase to show that DNA methylation is the primary determinant of MeCP2 binding in mouse brain. *In vitro* and *in vivo* analyses reveal that MeCP2 binding to non-CG methylated sites in brain is largely confined to the tri-nucleotide sequence mCAC. Structural modeling suggests that mCG and mCAC may be interchangeable as minimal structural perturbation of MeCP2 accompanies binding. MeCP2 binding to chromosomal DNA in mouse brain is proportional to mCG + mCAC density and defines domains within which transcription is sensitive to MeCP2 occupancy. The results suggest that MeCP2 interprets patterns of mCAC and mCG in the brain to negatively modulate transcription of genes critical for neuronal function.

## Introduction

Methylation at the C5 position of cytosine is an epigenetic mark implicated in gene regulation and disease^1^. In mammals, DNA methylation occurs in a CG di-nucleotide context, but in neuronal cells and embryonic stem cells (ESCs), mCA is detected at significant levels^2,3^. Like mCG, mCA is negatively correlated with transcript abundance, hinting at a repressive function in the brain^2,3^. Highest levels of mCA are observed in the human and mouse brain, where mCA accumulates postnatally during a phase of increased synaptogenesis^3^. In mice, the postnatal increase in neuronal mCA coincides with accumulation of Dnmt3a protein^3^. All cytosine DNA methyltransferases exhibit a preference for CG as a substrate, but Dnmt3a is able to methylate CA, albeit at a lower rate^4–7^. Brain mCA occurs most frequently in the tri-nucleotide mCAC^2,8^, whereas in ESCs mCAG is the preferred sequence context. The role of mCAC^2,3^ in the developing mammalian brain is yet to be elucidated.

A potential mechanism for interpreting the DNA methylation signal is the recruitment of methyl-CG binding domain (MBD) proteins including MeCP2, MBD1, MBD2 and MBD4^9^. Of these MeCP2 has attracted considerable attention as mutations involving the *MECP2* gene cause the X-linked autism spectrum disorder Rett syndrome^10^ and *MECP2* duplication syndrome^11^. Rett missense mutations cluster in two domains of MeCP2: the MBD and the NCoR/SMRT co-repressor interaction domain (NID)^12,13^. These observations implicate the loss of binding to methylated DNA and/or the failure to recruit the NCoR/SMRT repressor complex to DNA as primary causes of Rett syndrome. MeCP2 has a high affinity *in vivo* and *in vitro* for binding to mCG^14–16^, but the determinants of its targeting to DNA have recently diversified to include mCA, whose postnatal accumulation is paralleled by an increase in MeCP2 protein^2,3^. In addition, it has been reported that MeCP2 binds to hydroxymethylcytosine (hmC), the major oxidized form of mC, which is abundant in neurons^17^. Finally, there is data suggesting that MeCP2 can bind chromatin in a DNA methylation-independent manner^15,18,19^.

Although the mutational spectrum and biochemical interactions of MeCP2 suggest that it behaves as a transcription repressor^13,20^, changes in the mouse brain transcriptome when the protein is absent involve both up- and down-regulation of genes^17,21,22^. Accordingly MeCP2 has been proposed to act additionally as an activator of transcription or as a multifunctional hub that effects diverse aspects of cellular metabolism^12,23,24^. An alternative model proposes that MeCP2 primarily functions by globally modifying the architecture of chromatin via multifaceted interactions with DNA^16,19^. An inverse correlation between levels of CA methylation and expression of long genes has re-emphasized transcriptional inhibition via MeCP2^25^ A separate study reported, however, that mCA is enriched within genes that are misregulated regardless of the direction of the transcriptional change in response to MeCP2 depletion or excess^26^. Despite progress, therefore, a consensus view regarding the role of MeCP2 in transcriptional regulation has been elusive.

Here we define the DNA binding specificity of MeCP2 using *in vitro* and *in vivo* approaches. Supporting the primacy of DNA methylation as a determinant of MeCP2 binding, we find that global reduction of this epigenetic mark in the mouse brain decreases MeCP2 binding. We show for the first time that MeCP2 binding to non-CG methylated sites is primarily restricted to the tri-nucleotides mCAC or hmCAC *in vitro* and *in vivo.* Modeling based on the X-ray structure of the MBD of MeCP2 suggests that mCG, mCAC and hmCAC interact with a common protein conformation and may therefore lead to indistinguishable down-stream biological effects. MeCP2 binding across the adult brain genome reveals long genomic domains of high and low occupancy that match the distribution of mCG + mCAC binding sites. The results have important implications for regulation of gene expression, as we uncover a strong correlation between MeCP2 binding, mCG + mCAC density and the direction of gene mis-regulation when MeCP2 is absent or over-expressed. Our findings shed new light on the binding properties of MeCP2 and implicate MeCP2 as a global modulator of neuronal transcription.

## Results

### Global reduction of DNA methylation in brain causes loss of MeCP2 binding

DNA methylation as a determinant of MeCP2 binding has been demonstrated experimentally^15,16,27^, but its primary role has been questioned^28^ and the possibility considered that MeCP2 binds to DNA via multiple modalities^18^. Depletion of the CG methylation signal specifically in brain would stringently test its importance, but mice in which *Dnmt1* is deleted exclusively in neuronal and glial cells using Nestin–Cre die at birth^29^. Reinvestigating this conditional mutation strategy, we found that on an outbred background mice survive postnatally for up to two weeks. At this age, neuronal MeCP2 expression levels are low compared to adult brains^30^ and MeCP2 gains in functional importance only later in development as *Mecp2* knockout (KO) mice are symptom-free at this stage^31,32^. Nevertheless, *Dnmt1* KO mice represent a unique opportunity to test for causal effects of DNA methylation on MeCP2 binding *in vivo.*

Western blots confirmed that Dnmt1 was severely reduced from birth to postnatal day 13 in brains expressing Nestin–Cre and the floxed *Dnmt1* allele (Fig. 1a). Moreover transcription of intracisternal A particle *(IAP)* elements, which are members of an LTR retrotransposon family in mouse, was greatly up-regulated as reported in Dnmt1-deficient mouse embryos^33^ (Fig. 1b), but expression of other transposon families was not detectably affected (Supplementary Fig. 1a-b). *Mecp2* -/y brains showed no significant elevation of *IAP* element transcription (Fig. 1b). A previous study^16^ found that *IAP* transcription was modestly increased (∼ 2-fold) in nuclear RNA from *Mecp2*-null brain, but this effect was not detectable in stable RNA from whole tissue, in agreement with present findings. We analyzed levels of global DNA methylation in wildtype (WT) and *Dnmt1* KO brains by high performance liquid chromatography (HPLC) (Fig. 1c and Supplementary Fig. 1c) and reduced representation bisulfite sequencing^34^ (RRBS) (Fig. 1d and Supplementary Fig. 1d). The unbiased HPLC quantification detected a 38% reduction in CG methylation in *Dnmt1* KO brains whereas the RRBS tested regions, which comprise predominantly promoter regions, showed a 56% reduction. Importantly, MeCP2 was expressed at near normal protein levels in *Dnmt1* KO brains in comparison with age-matched WT brains (Fig. 1e-f and Supplementary Fig. 1e).

**Figure 1.**
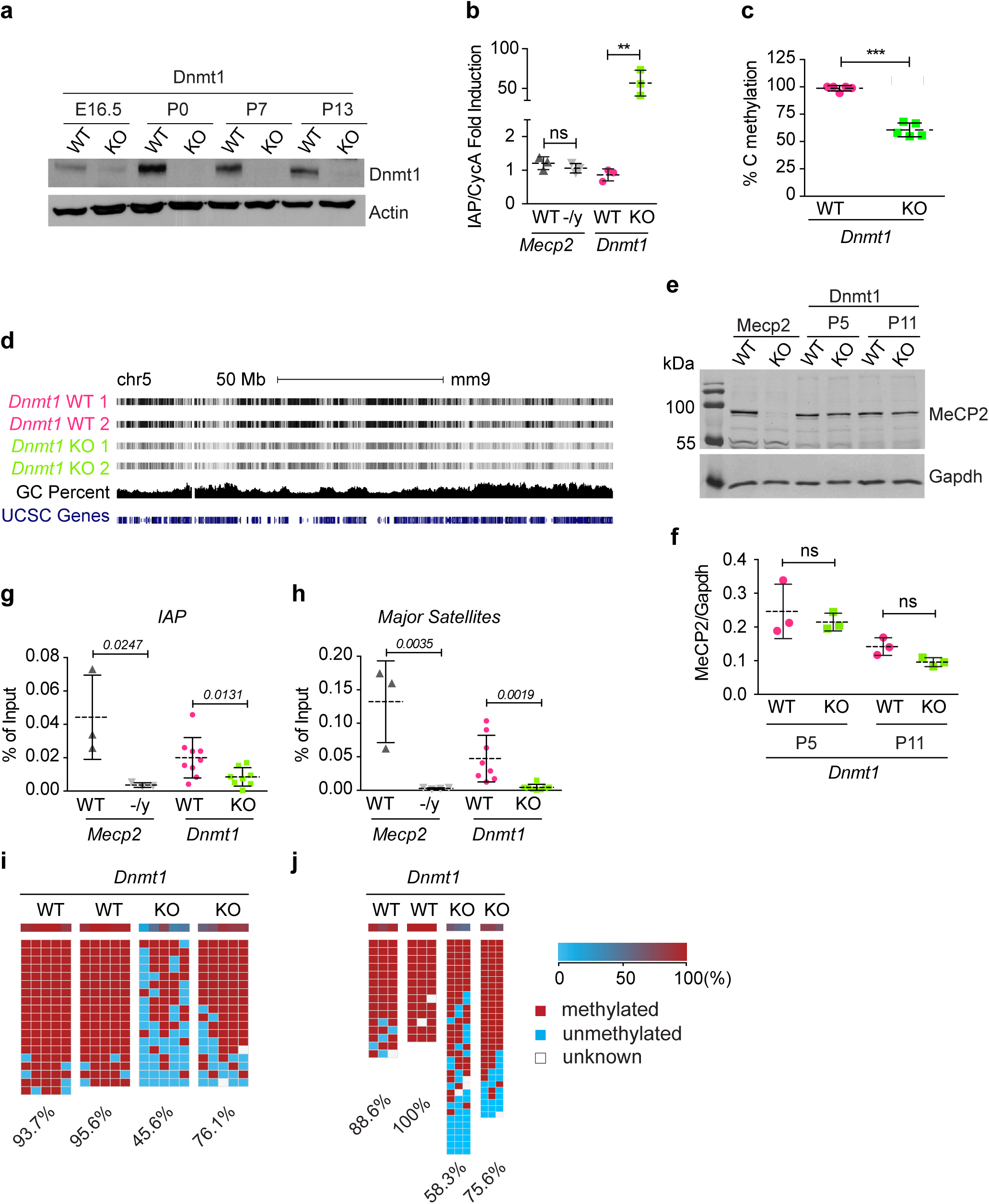
DNA methylation is the primary determinant of MeCP2 binding *in vivo.* (a) Western blot of whole brain from WT and *Dnmt1* conditional KO mice at different age points (E = embryonic day, P = postnatal day) with antibodies against Dnmt1 and β-Actin. (b) Real Time PCR of *IAP* expression in WT and *Mecp2* -/y male (8 weeks old) and WT and *Dnmt1* KO (P11) triplicate whole brain cDNAs normalized to *Cyclophilin A. Mecp2* WT and *Dnmt1* WT data points were set to 1. (c) HPLC analysis of % total C methylation in 5 samples of P12 and P5 WT and *Dnmt1* KO whole brain. WT methylation levels set to 100%. Individual replicates at two age points are shown in Supplementary Fig. 1c. (d) RRBS of WT and *Dnmt1* KO brain duplicates at P5. UCSC browser screenshot of chromosome 5. Scale bar = 50 Mb. Grey levels indicate methylation levels between 0 (white) and 100% (black). (e) Western blot of WT, *Mecp2* KO adult and WT, *Dnmt1* KO whole brain at P5 and P11 with antibodies against MeCP2 and Gapdh. (f) Quantification of triplicate WT and *Dnmt1* KO whole brain at P5 and P11. Western blot for quantification is shown in Supplementary Fig. 1e. (g-h) ChIP of WT and *Mecp2* KO adult and WT and *Dnmt1* KO replicates at *IAP* (g) and major satellite repeats (h) at P12. Results are presented as % of input. All error bars represent ± SD. P values were calculated with Students unpaired one-tailed t-test (g-h) or Students unpaired two-tailed t-test (b-c, f): ns p>0.05; * p<0.05; ** p<0.01, ***p<0.001. (i-j) Bisulfite sequencing 1 of *IAP* (i) and major satellite repeats (j) in WT and *Dnmt1* KO genomic DNA from whole brain at P12. Percentage of methylation is indicated. Rows correspond to individual sequenced DNA strands.

To investigate how local loss of DNA methylation quantitatively affects MeCP2 binding by chromatin IP (ChIP), we used primers specific for repetitive *IAP* elements and mouse major satellite sequences (Fig. 1g-h and Supplementary Fig. 1f) plus four inter- and intra-genic single-copy regions from mouse chromosome 3 (Supplementary Fig. 1g-k). Reduced DNA methylation in the conditional mutant brain was confirmed at *IAP* elements and mouse satellite by bisulfite analysis (Fig. 1i-j) and MeCP2 binding was correspondingly lower (Fig. 1g-h). Extending the analysis to unique regions of the genome, we chose four random sites in a relatively gene dense domain (Supplementary Fig. 1g). In all replicates MeCP2 binding to each site was consistently reduced in each *Dnmt1*-deficient brain compared with a matched WT littermate (Supplementary Fig. 1h-k). Furthermore, bisulfite analysis confirmed the reduction of DNA methylation at the nucleotide level in each case (Supplementary Fig. 1l-o). The data for 6 randomly chosen regions of the mouse genome provide strong support for the conclusion that DNA methylation is a major determinant of MeCP2 binding in the mouse brain.

### MeCP2 recognizes modified di- and tri-nucleotide sequences

The predominant methylated sequence is the di-nucleotide CG, but in adult brain mCA^3^ and hmCG^17,35^ are implicated as binding partners of MeCP2. A recent study concluded that in addition to mCG, MeCP2 binds both mCA and hmCA^25^ and confirmed earlier reports that hmCG is a low-affinity binding site for MeCP2^25,36-38^. In addition MeCP2 has been reported to bind *in vitro* to DNA in which every cytosine was substituted with hmC^17^. Given that neurons only accumulate significant levels of both hmCG and mCH (where H is any base except G) from two weeks postnatally^3^, the *Dnmt1* KO model is unsuited to investigate additional sequence specificities of methylation dependent MeCP2 binding. To comprehensively analyze the DNA sequence determinants of MeCP2 binding *in vitro* we performed EMSA using the 1-205 N-terminal domain of MeCP2 that contains the MBD. Surprisingly, the data revealed a further constraint on MeCP2 binding, as the third base following mCA strongly affected MeCP2 binding affinity *in vitro* (Fig. 2a). Probes containing the mCAC tri-nucleotide sequence bound with high affinity to MeCP2, whereas probes containing mCAA, mCAG and mCAT bound much less strongly. This result was confirmed in EMSA experiments using all possible mCXX tri-nucleotide sequences as unlabeled competitors against a labeled mCGG-containing probe (Fig. 2b). Quantification showed that mCAC and, to a lesser extent mCAT, are both effective competitors, but mCAG and mCAA compete no better than non-methylated control DNA (Fig. 2c). All mCGX oligonucleotide duplexes competed strongly indicating that the base following mCG on the 3’ side does not have a large effect on binding, although we note that mCGA was reproducibly a weaker competitor than mCGC, mCGG or mCGT. Neither mCCX nor mCTX tri-nucleotides have a significant affinity for MeCP2 *in vitro.*

**Figure 2.**
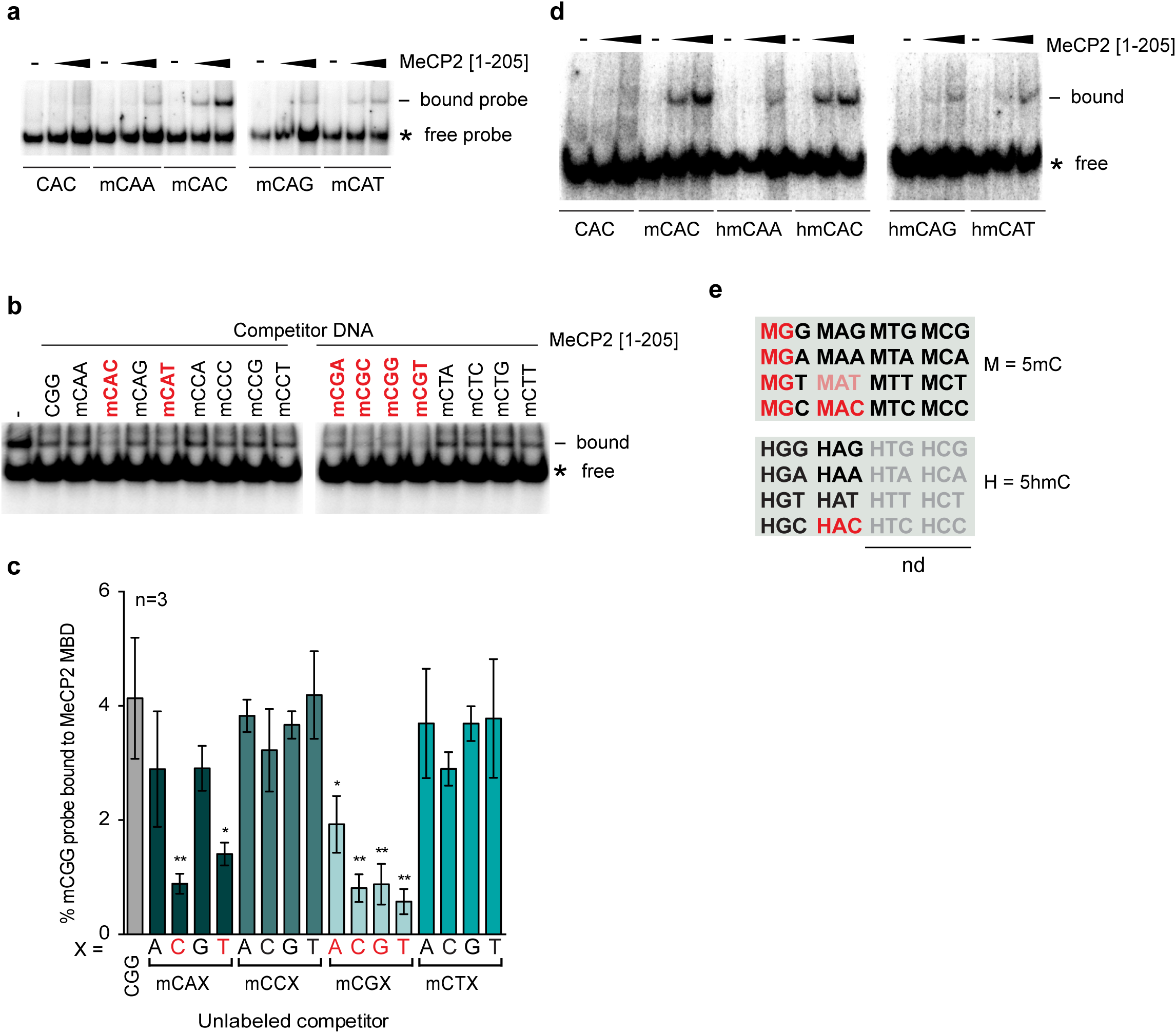
MeCP2 binds mCAC and hmCAC *in vitro.* (a) EMSA using no protein (-) or varying amounts of MeCP2 [1-205] with a probe (Supplementary Table 1) containing a centrally methylated C in a CAX context. Gap indicates separate gels. (b) EMSA using MeCP2 [1-205] or no added protein (-) in the presence of excess unlabeled unmethylated competitor DNA (CGG) or methylated competitor DNA (mCXX). Labeled probe contains mCGG. Red denotes strongest competition. (c) Quantification of (b). Three individual experiments were averaged. Red denotes most significant competition. Significance was calculated in relation to unmethylated CGG (grey bar). Error bars represent ± SD. Students unpaired t-test: * p<0.05; ** p<0.01. (d) EMSA using no protein (-) or varying amounts of MeCP2 [1-205] with a probe containing a centrally hydroxymethylated C in a CAX triplet context. * : free probe; - : bound probe. (e) Summary of MeCP2 binding motifs *in vitro.* M = 5-methylcytosine, H = 5-hydroxymethylcytosine. Bright red: strong binding; pale red: weaker binding; grey: not tested.

As hmCA is reported to bind MeCP2 *in vitro^25^,* we asked whether the third base is also important for hmC binding. Using hmCXX tri-nucleotides as probes in EMSAs, we found that hmCAC bound with a much higher affinity than hmCAA, hmCAG and hmCAT DNA (Fig. 2d). It is notable that the great majority of hmC in the brain and elsewhere is in the hmCG di-nucleotide, with hmCAC being extremely rare^3^. The DNA binding specificity of MeCP2 MBD deduced from these *in vitro* experiments is summarized in a matrix of di- and tri-nucleotide sequences that bind to MeCP2 (red lettering, Fig. 2e).

### MeCP2 binding specificity in vivo

To determine whether the binding specificities established *in vitro* apply to full-length protein in living cells, we developed a novel assay using transfection followed by ChIP^39^. Synthetic DNA duplexes containing specific cytosine modifications were transfected into HEK293 cells expressing human MeCP2 tagged with GFP (Fig. 3a and Supplementary Fig. 2a-b). Levels of endogenous MeCP2 in these cells are negligible and therefore do not interfere with the assay (Supplementary Fig. 2c). We tested oligonucleotide duplexes containing a single modified cytosine in either a mCG, hmCG, mCAX or hmCAX context (Fig. 3b). The results showed that mCAC, mCAT, hmCAC and mCG all bound MeCP2 efficiently, whereas hmCG, mCAA and mCAG binding was indistinguishable from background binding to non-methylated DNA (Fig. 3c). The same outcome was seen with different DNA sequences containing three CAC motifs per oligonucleotide, either all unmodified, all methylated or all hydroxymethylated (Fig. 3d-e). These *in vivo* results with full-length protein match those obtained *in vitro* by EMSA, strengthening the evidence that MeCP2 requires specific tri-nucleotide settings to recognize mC or hmC in a non-CG context.

**Figure 3.**
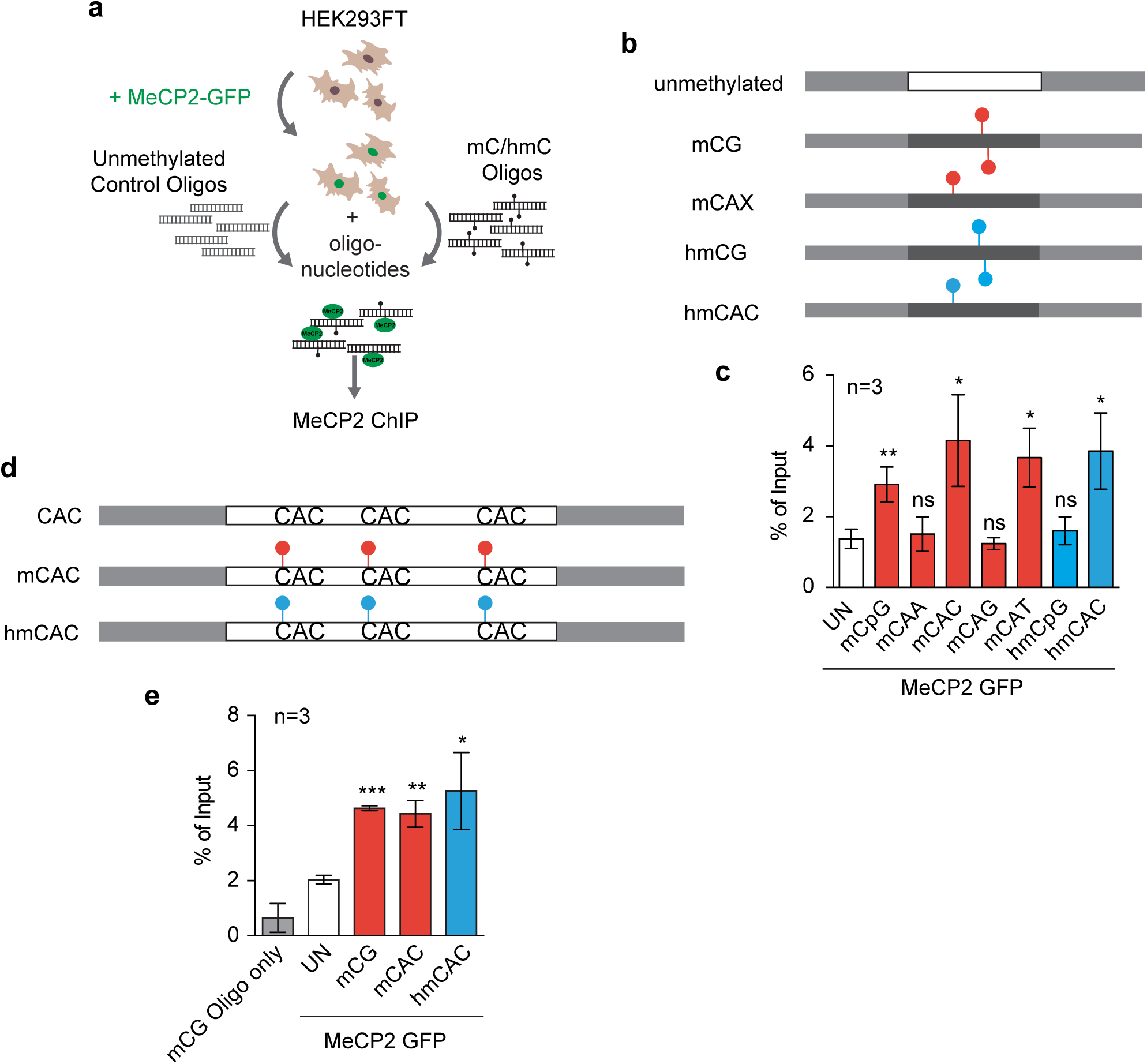
Full-length MeCP2 binds mCAC and hmCAC *in vivo.* (a) Schematic of *in vivo* transfection assay in HEK293FT cells. (b) Differentially modified oligonucleotide derived from the mouse *Bdnf* locus. Light grey: T3 and M13-20 adapters; red circles: mC; blue circles: hmC. See also Supplementary Table 1. (c) Real Time PCR of *in vivo* transfection assay in triplicate where WT MeCP2-GFP was co-transfected with oligonucleotide as described in (b). (d) Differentially modified oligonucleotide at three CAC sites. See also Supplementary Table 1. Light grey: T3 and M13-20 adapters; red circles: mC; blue circles: hmC. (e) Real Time PCR of *in vivo* transfection assay in triplicates where WT MeCP2-GFP was co-transfected with oligonucleotides as described in (d). Real Time PCR results are presented as *%* of Input (red bars: mC; blue bars: hmC, white bars: unmethylated; grey bar: mCG oligonucleotide transfected without prior transfection of MeCP2-GFP as a background control). Error bars represent ± SD. Significance was calculated in relation to unmethylated oligonucleotide transfections (white bars). Students unpaired t-test: ns p>0.05; * p<0.05; ** p<0.01; *** p<0.001.

### Modeling the MeCP2-DNA interaction

To seek a structural basis for the tri-nucleotide specificity of MeCP2 binding, we asked whether the X-ray structure of the MeCP2 MBD^40^ could suggest why mCAC or mCAT binding is permitted while mCAA, mCAG, mCCX and mCTX are forbidden. Surprisingly, altering the conformation of only one amino acid side chain, R133, while leaving all other coordinates of the protein structure unchanged, could account for the observed interactions. R133 makes essential hydrogen bonds with one guanine base in the mCG complex and also provides salt bridges with a cytosine methyl group (Fig. 4a)^40^. The equivalent guanine residue on the other DNA strand of the mCG dyad is also hydrogen bonded to an arginine: residue R111. Mutations in either R133 or R111 cause Rett syndrome, but despite their related roles, the conformations of R111 and R133 are very different. Whereas the R111 side-chain adopts an extended all-trans conformation that is “pinned” by hydrogen bonds with D121, R133 is relatively unconstrained by surrounding amino acids (Fig. 4a). We therefore asked if R133 could potentially interact with mCAC in the existing X-ray structure^40^. Indeed, by extending the side-chain and tilting the guanidinium group, R133 can make permitted stabilizing hydrogen bonds with the guanine base that is paired with the third cytosine in the mCAC tri-nucleotide (Fig. 4b). An equivalent interaction is also possible with the complementary adenine in mCAT tri-nucleotide (Supplementary Fig. 3a). Importantly, interactions with pyrimidine bases are sterically unfavorable, arguing that mCAA or mCAG are unlikely to interact with the MBD, as is observed experimentally (Supplementary Fig. 3b-c). Modeling of hmCG binding indicates that the presence of the cytosine hydroxyl group would not be accommodated due to the close proximity of this polar group with the guanidinium group of R111 (Supplementary Fig. 3d). Binding of hmCAC is allowed, however, due to tilting of the R133 side-chain and the formation of hydrogen bonds with guanine on the opposite DNA strand (Supplementary Fig. 3e). The presence of a purine in the third position of hmCAG and hmCAA introduces a clash that is not observed with hmCAC or hmCAT (data not shown). Thus the observed tri-nucleotide binding specificity of MeCP2 can theoretically be explained with minimal perturbation of the established X-ray structure of the MBD-DNA complex. The minimal change to the structure of the MBD required to accommodate either mCG or mCAC raises the possibility that the biological consequences of binding either motif will be the same.

**Figure 4.**
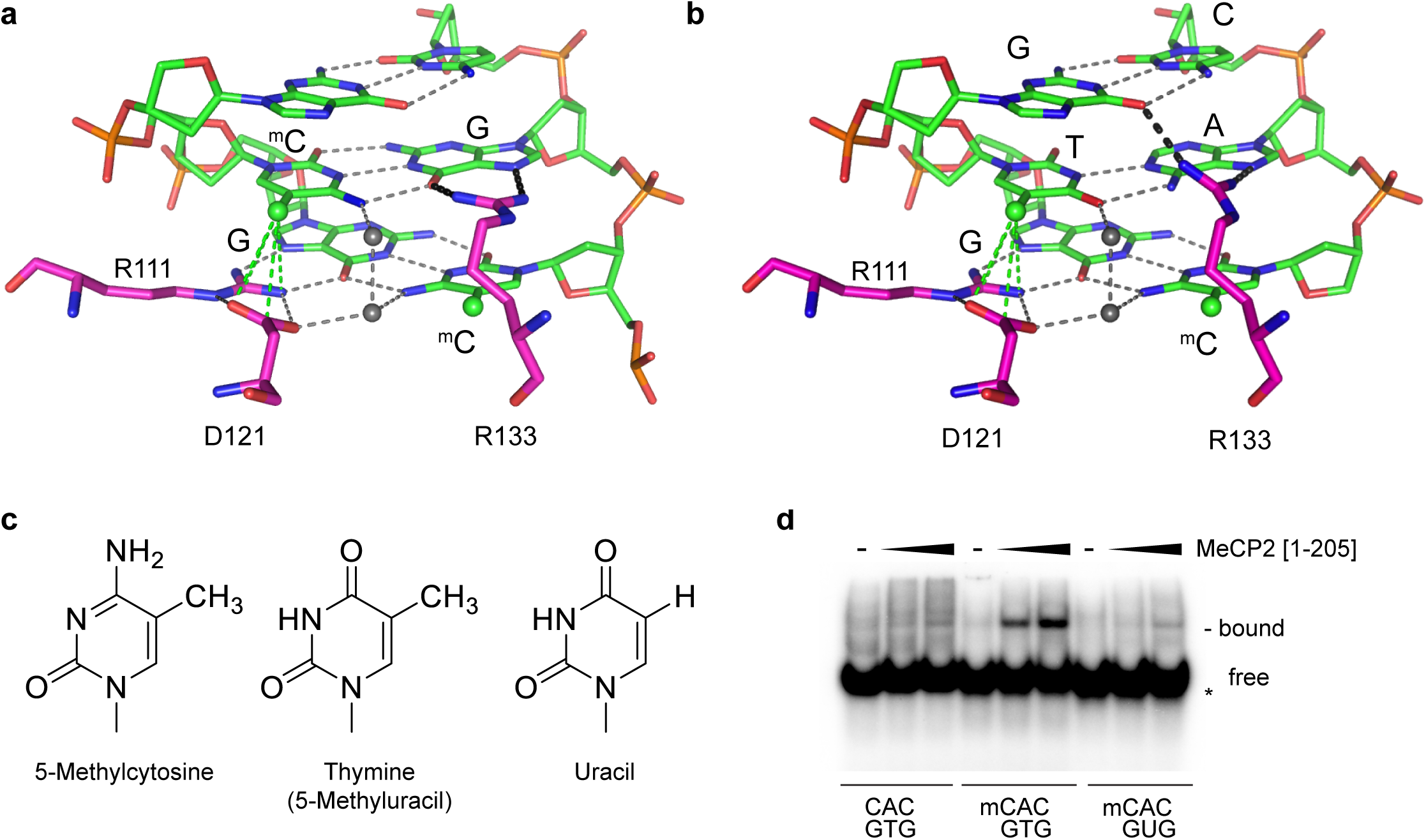
MeCP2 tri-nucleotide recognition may depend on flexibility of the R133 side chain and requires the methyl group of thymine. (a) X-ray structure of MeCP2 [77-167] interaction with the mCG di-nucleotide in double-strand DNA^40^. Critical amino acids R111, R133 and D121 are shown in pink. Green sphere: methyl group; grey sphere: water molecule; green dotted lines (from R111 and D121): favorable interactions; grey dotted lines: hydrogen bonds. (b) Modeling of the MeCP2 [77-167] interaction with double-strand DNA containing mCAC. Key as in (a) and black dotted lines: modeled interactions. (c) Structure of 5-methylcytosine, thymine and uracil. Note that thymine and uracil are distinguishable by a methyl group on position 5 of the pyrimidine ring. (d) EMSA using no (-) or varying amounts of MeCP2 [1-205] to assess the influence of the methyl group of thymine on binding. * : free probe; - : bound probe.

A prediction of the modeling is that MeCP2 should bind in only one orientation to mCAC or mCAT, whereas binding to the symmetrical mCG dyad may occur in either orientation. The model requires that R111 and D121 interact with the methyl group of thymine rather than that of 5mC (Fig. 4b). As thymine is effectively 5-methyluracil (Fig. 4c), we replaced it with uracil in the labeled probe and performed EMSA analysis (Fig. 4d). Loss of the thymine methyl group abolished binding to MeCP2, in agreement with the hypothesis that MeCP2 binding to mCAC is confined to one orientation. The data suggest that a symmetrical pair of 5-methyl pyrimidines, one of which is mC, offset by one base pair is an essential pre-requisite for MBD binding to DNA.

### DNA sequence specificity of MeCP2 binding in adult mouse brain

To test whether the DNA binding specificities established *in vitro* and in transfected cells also apply in native tissues, we analyzed MeCP2 ChIP-seq and whole genome bisulfite (WGBS) datasets derived from adult mouse brain (references ^3,25,26^ and WGBS from sorted neurons and WGBS from hypothalamus this study). CG is under-represented in the mouse genome (∼4% of CX), but highly methylated (∼80%), whereas CA is the most abundant CX di-nucleotide (36% of CX), but even in brain only a small fraction of CA is methylated (<2%) (Fig. 5a-c). Bisulfite analysis of sorted neurons by NeuN staining^16^ confirmed previous reports that CAC is the most methylated tri-nucleotide^2,8^, being ∼12% methylated (Fig. 5c). The finding that the MeCP2 tri-nucleotide binding specificity matches the most abundantly methylated non-CG sequence in brain encourages the view that this interaction is biologically relevant.

**Figure 5.**
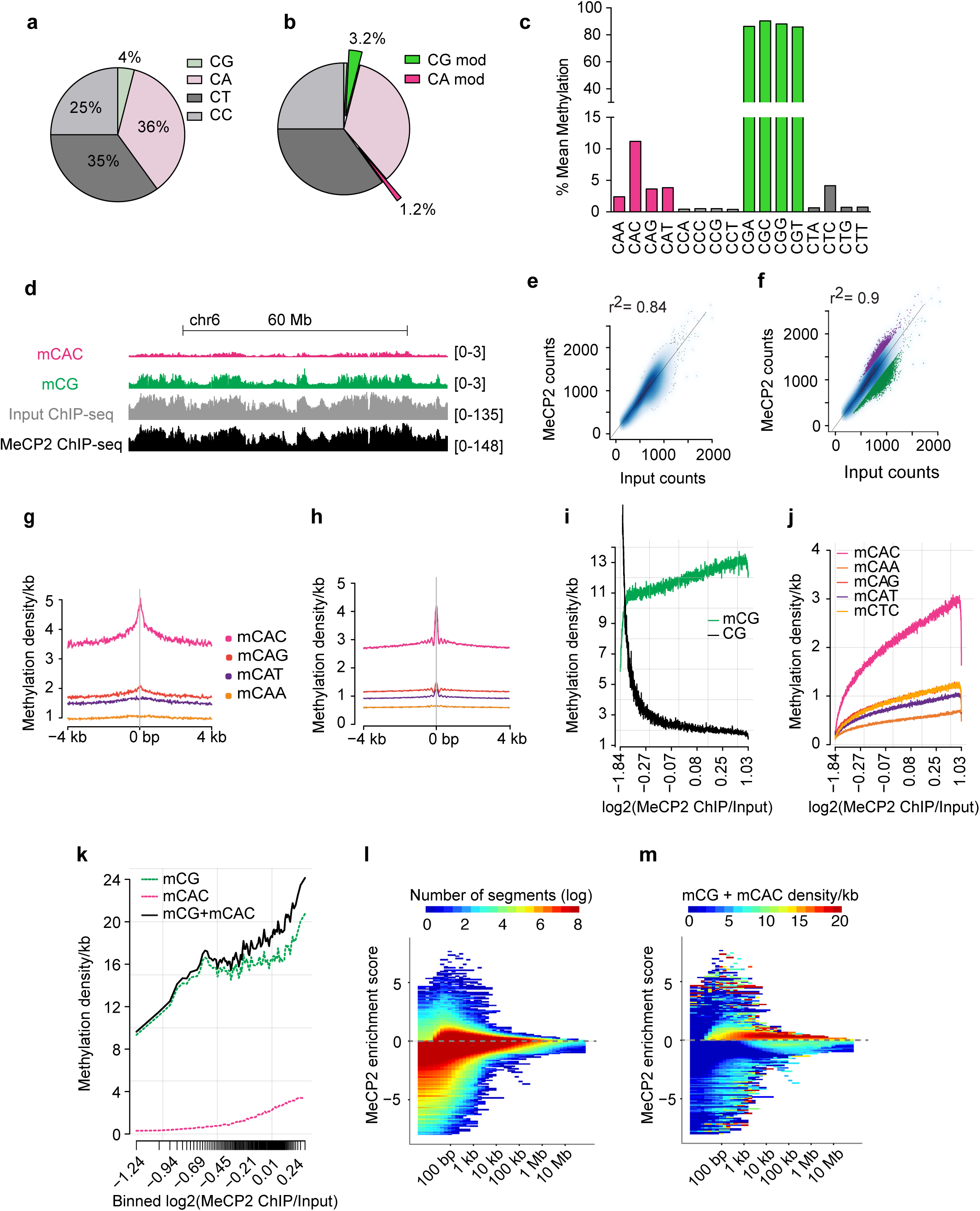
DNA methylation and MeCP2 binding in the mouse brain. (a) Pie chart showing CX frequencies in the mouse genome (dataset WGBS of sorted neurons from this study). (b) Pie chart showing modified (mCG and hmCG) CX frequencies in brain neuronal nuclei as determined by WGBS (dataset WGBS of sorted neurons from this study). (c) Mean methylation levels (%) of CXX in brain neuronal nuclei based on WGBS of sorted neurons (dataset from this study). (d) IGV browser screenshot of mCAC, mCG, MeCP2 ChIP-seq and corresponding Input DNA based on sequence reads. Data represent a 100 Mb region of chromosome 6 (datasets from ^3,26^). (e) Correlation between sequence coverage of MeCP2 ChIP-seq and corresponding Input DNA in 10 kb windows. Fitting a linear model (MeCP2 ∼ Input) yields a coefficient of determination of 0.84 (dataset from ^26^). (f) As for (e) but highlighting relative depletion (4.1% of windows; green) and relative enrichment (1.7% of windows; purple). When these outliers are excluded, 90% of the variability of the MeCP2 signal in the remaining binned windows is explained by sequence bias (Input coverage) (dataset from ^26^). (g-h) Enrichment of mCAC at summits of MeCP2 binding (datasets from ^3,25^ (g) and ^26^ and hypothalamus WGBS from this study (h)). (i) Relationship between mCG and unmethylated CG density/kb and MeCP2 occupancy corrected for Input in genome-wide 1 kb windows (datasets from ^26^ and hypothalamus WGBS from this study). (j) Relationship between mCAX and mCTC density/kb and MeCP2 occupancy corrected for Input in genome-wide 1 kb windows (datasets from ^26^ and hypothalamus WGBS from this study). (k) Methylation density as a function of log2(MeCP2 ChIP/Input) in protein-coding genes. The density of tick marks on the x-axis represents the distribution of genes with respect to MeCP2 binding (datasets from ^26^ and hypothalamus WGBS from this study). (l-m) Domains of MeCP2 enrichment and depletion as determined by MSR (datasets from ^26^ and hypothalamus WGBS from this study). Grey dotted line indicates zero. (l) Heatmap showing number of segments binned by their scale (x-axis) and MeCP2 enrichment scores (y-axis). X-axis shows the median lengths of segments found with a given scale. Positive scores on the y-axis indicate MeCP2 enrichment; negative scores indicate MeCP2 depletion. Plot is colored according to number of segments. (m) Heatmap showing number of segments binned by scale and scored as in (l). Plot is colored according to combined mCG + mCAC density.

Previous ChIP-seq analyses have concluded that the read coverage of MeCP2 chipped samples tracks the density of CG methylation^15,16,25,26^ Reanalysis of several MeCP2-ChIP data sets for which the antibody used has been rigorously verified, indicates, however, that the profile of Input – specifically, DNA derived from the fragmented chromatin sample used for ChIP – is closely similar to that of MeCP2 (Fig. 5d). Using the published high quality ChIP-seq dataset for hypothalamus^26^, we fitted a linear model to predict MeCP2 read coverage from Input reads alone and found a coefficient of determination of 0.84, indicating that MeCP2 is almost uniformly distributed across the genome at this resolution (Fig. 5e). These results are in line with a previous report that the number of MeCP2 molecules in mature neurons is sufficient to almost ‘‘saturate’’ mCG sites in the genome^16^. Given the similarity between ChIP and Input, we focussed on regions that show deviations from the Input profile regarding enrichment (purple) or depletion (green) of MeCP2 (Fig. 5f). First, we investigated genomic regions that are depleted of potential binding sites, e.g. unmethylated CpG islands (CGIs). As the ChIP dataset was derived from mouse hypothalamus, we performed WGBS on three biological replicates of this brain region (see Online Methods). Using ChIP and DNA methylation datasets from the same brain region, we observed a pronounced drop of the log2(MeCP2/Input) signal across CGIs for two different data sets, confirming previous analyses^16,25,26^ (Supplementary Fig. 4a). We next examined regions where the MeCP2 signal was higher than expected by applying the MACS^41^ tool to detect summits of MeCP2 binding peaks relative to Input^25^. As expected, the di-nucleotide mCG showed a sharp peak at MeCP2 ChIP summits in the hypothalamus dataset (Supplementary Fig. 4b-e and ^25^). In addition, the tri-nucleotide mCAC, but not other mCAX tri-nucleotides, coincided strikingly with MeCP2 peak summits, confirming that mCAC provides a focus for MeCP2 binding (Fig. 5g-h and Supplementary Fig. 4f-h). Using random regions as a negative control, we did not detect any sequence or methylation dependency (Supplementary Fig. 4i). Regarding targeting of mCAT, which bound relatively weakly in EMSA, but strongly in the transfection assay, the ChIP-seq data suggest that this is a relatively low affinity site in native brain (Fig. 5g-h and Supplementary Fig. 4f-h).

To establish MeCP2 binding preferences across the whole genome we adopted a sliding window approach. In the Input sample, both, mCG density and density of unmethylated CGs strongly correlated with coverage revealing a methylation independent CG sequencing bias (data not shown). In contrast, the MeCP2 ChIP coverage was DNA methylation sensitive, with mCG being positively correlated, while unmethylated CG density was anti-correlated (data not shown). Importantly, DNA methylation-sensitivity was also observed in the Input-corrected signal (log2(MeCP2/Input)), strongly supporting the view that mCG is targeted by MeCP2 binding (Fig. 5i, Supplementary Fig. 4j, l). This mCG binding preference was independent of the third DNA base (Supplementary Fig. 4n). In agreement with the *in vitro* and *in vivo* results reported above, the density of mCAC correlated strongly with increasing MeCP2 enrichment, whereas a much weaker trend was observed for other methylated tri-nucleotide sequences (Fig. 5j, Supplementary Fig. 4k, m). We found no evidence for MeCP2 binding to hmCG or unmethylated C’s in any sequence context (Supplementary Fig. 4j, l, o-r). To complement the sliding window approach, we focused on MeCP2 binding within gene bodies and found once again that the Input-corrected signal is strongly correlated with the density of mCG and mCAC (Fig. 5k). The ChIP data therefore sustain the view that MeCP2 binding is determined by the combined density of mCG and mCAC sites.

Given the high abundance and global distribution of MeCP2 in the neuronal genome, we looked for domains of MeCP2 occupancy that might reflect long-range variation in binding site abundance. To avoid pooling data in arbitrary windows, we used a multiscale representation method (MSR) that identifies patterns of signal enrichment or depletion across scales spanning several orders of magnitude^42^. MSR identified a large number of long domains moderately enriched for MeCP2 extending up to 1 Mb indicating that regions of high MeCP2 occupancy extend beyond the scale of a single gene (Fig. 5l). These regions share common sequence features, in particular high mCG and mCAC densities (Fig. 5m, Supplementary Fig. 4t-u). We also identified a large number of short regions containing highest GC content (<1 kb) but are strongly depleted in MeCP2 binding. As expected, these regions significantly overlap with CGIs (Supplementary Fig. 4s). Lastly, we found a third group of relatively long regions (10 kb - 1 Mb) which are moderately depleted in MeCP2 binding (MeCP2 enrichment score: <0). These regions are moderately enriched for mCG but lack mCAC (Fig. 5m, Supplementary Fig. 4s-u).

As part of this binding site analysis we re-visited an earlier report that MeCP2 binds preferentially to mCG flanked by an AT-rich run of 4-6 base pairs *in vitro*^43^. To look for this preference in brain, we asked whether isolated mCG and mCAC flanked by a run of 4 or more A or T base pairs within 13 base pairs showed greater MeCP2 ChIP-seq signal than sequences lacking an AT run. In summary, we find no evidence for an effect of AT-flanks on MeCP2 binding site occupancy, suggesting that the *in vitro* preference might not be biologically relevant *in vivo* (Supplementary Fig. 4v-w).

### The relationship between MeCP2 occupancy and gene expression

The association of MeCP2 with both methylated sites in the genome and the co-repressor complex NCoR^13^ suggests that the protein can function to inhibit transcription. If so, a relationship would be expected between MeCP2 occupancy and the transcriptional mis-regulation seen when MeCP2 is either depleted by deletion of the gene (KO)^31^ or over-expressed (OE)^44,45^. Before making use of published datasets for mouse hypothalamus^26^, we first asked, using HPLC, whether the absence or over-expression of MeCP2 alters total RNA levels. Previous studies using MeCP2-deficient neurons differentiated *in vitro* from mouse ES cells or human iPS cells reported reduced total RNA and transcriptional capacity^24,46^, but comparable measurements in brains of MeCP2-deficient mice have not been reported. Using a sensitive RNA quantification technique, we observed that total RNA per cell in KO hypothalamus is reproducibly 15% lower than WT (Supplementary Fig. 5a). Over-expression of MeCP2, however, did not significantly affect total RNA. As the latter is ∼98% ribosomal RNA, we asked whether mRNA levels were also reduced in KO hypothalamus. Quantitative RT-PCR (qPCR) using spiked-in *Drosophila* cells to control for experimental error and normalized to brain cell number in each sample (Supplementary Fig. 5b) confirmed that genes previously reported to be up- or down-regulated in MeCP2-deficient hypothalamus^26^ were similarly mis-regulated in our samples (Supplementary Fig. 5c-f). We then measured the abundance of three housekeeping genes and found that all were down-regulated by approximately 15% (Supplementary Fig. 5g-i). These results suggest that total RNA and mRNA levels are coordinately reduced and we have therefore applied this normalization to all hypothalamus RNAseq datasets. The mechanisms responsible for reduced total RNA, and whether this effect is a primary or secondary consequence of MeCP2 deficiency, are unknown.

We next examined gene expression by separating genes whose expression was increased, unchanged or decreased in response to changing levels of MeCP2. Normalizing ChIP signals against Input, we found that up-regulated genes in KO hypothalamus are within domains enriched in MeCP2, whereas down-regulated genes were relatively depleted (Fig. 6a). Unchanged genes showed an intermediate level of MeCP2 occupancy. The reciprocal result was seen in OE hypothalamus, as down-regulated genes had high MeCP2 occupancy, whereas up-regulated and unchanged genes bound relatively less MeCP2 (Fig. 6d). This relationship, which was not observed in a previous analysis of this gene expression and ChIP data^26^, disappeared altogether if gene body binding of MeCP2 was normalized to binding in gene flanking regions, as enhanced or depleted binding extended up- and down-stream of the transcription start and end sites. We also asked whether the increased MeCP2 occupancy measured by ChIP-seq in hypothalamus correlated with an elevated level of the two target sequences mCAC and mCG. The distribution of mCAC strikingly matched the pattern of MeCP2 binding (Fig. 6b, e), but mCG, which occurs at much higher density, correlated less obviously (Fig. 6c, f).

**Figure 6.**
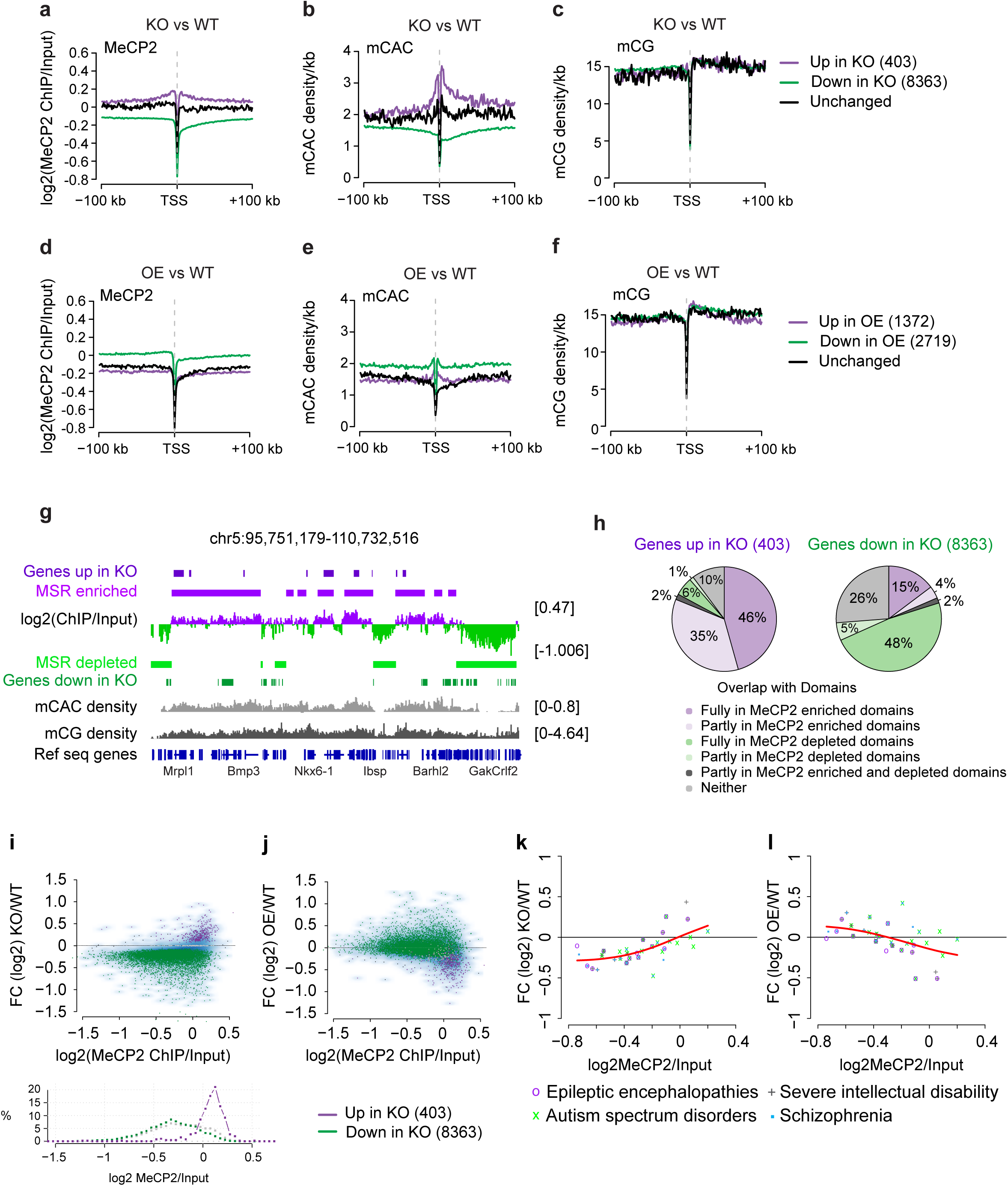
Genes show global down-regulation in KO, but MeCP2 enriched genes are up-regulated upon MeCP2 depletion and down-regulated when MeCP2 is overexpressed. Datasets ^26^ and hypothalamus WGBS from this study. (a, d) Aggregate WT MeCP2 occupancy plotted 100 kb up- and downstream of the transcription start site (TSS) of genes that are up-regulated (purple), down-regulated (green) or unchanged (black) in (a) *Mecp2* KO or (d) *Mecp2* OE hypothalamus. (b, e) Methylation density per kb of CAC and (c, f) CG plotted 100 kb up- and down-stream of the TSS using the same gene sets as in (a) and (d) respectively. (g) IGV screenshot of chromosome 5 (chr5:95,751,179-110,732,516) showing unpruned MSR regions of scale 30, corresponding to a median segment length of 270 kb and 240 kb for MeCP2 enriched and MeCP2 depleted segments. Datasets ^26^ and hypothalamus WGBS from this study. (h) Pie charts showing percentage of up-(left) and down-regulated (right) genes and their overlap with MeCP2 enriched (purple) or depleted (green) domains. Dataset ^26^ and hypothalamus WGBS from this study. (i) Transcriptional changes resulting from MeCP2 deficiency as a function of MeCP2 occupancy. Genes that are up-regulated in KO (purple) have high MeCP2 occupancy and show the strongest down-regulation in OE (j). FC = fold change. Histogram depicting the correlation between MeCP2 occupancy and *%* of genes that are up-(purple) or down-regulated (green). Dataset ^26^. (k-l) MeCP2 occupancy and expression changes are well correlated at genes previously implicated in neurological diseases^47^: Spearman correlation coefficients of 0.67 for KO vs WT (k) and -0.5 for OE vs WT (l) Dataset ^26^

The strong reciprocal relationship between MeCP2 occupancy and the direction of gene mis-regulation in KO and OE hypothalamus respectively, is compatible with the notion that MeCP2 binding is inhibitory to transcription. Excess MeCP2 preferentially inhibits genes with most binding sites whereas its depletion preferentially de-represses highly occupied genes. Elevated or depleted MeCP2 binding extended up- and down-stream of the TSS, suggesting that these genes are embedded within the extended MeCP2 domains identified by our MSR analysis (Fig. 5l-m). This was confirmed by mapping MeCP2-enriched and depleted domains onto the genome in relation to mis-regulated genes (Fig. 6g). Approximately 80% of genes up-regulated in KO hypothalamus were within or overlapped domains of high MeCP2 occupancy (dark and pale lilac, left pie), whereas only 19% of down-regulated genes were associated with MeCP2 enrichment (dark and pale lilac, right pie) (Fig. 6h and Supplementary Fig. 6a-b).

In order to comprehensively visualize the relationship between transcription and MeCP2 occupancy, all reliably detectable transcript levels were plotted against MeCP2 ChIP signal normalized to Input (Fig. 6i-j and Supplementary Fig. 6e-f). The 15% global reduction in mRNA and ribosomes meant that in KO hypothalamus expression of most genes (8,363) goes down (Fig. 6i, green), but despite this overall trend a small number of genes (403) significantly increased their expression per cell compared with WT (Fig. 6i, purple). These genes, which shared high MeCP2 occupancy (Fig. 6i, lower panel), showed the inverse behavior in OE hypothalamus where they were down-regulated (Fig. 6j). This inverse relationship is readily apparent in a plot of gene expression changes in KO/OE hypothalamus (Supplementary Fig. 6g). De-regulated genes shared similar levels of mCG density, but only the up-regulated genes displayed increased mCAC (Supplementary Fig. 6h-i). We noted that the genes up-regulated in KO were significantly longer than average, in agreement with a report that long genes are preferentially up-regulated in MeCP2 KO mice^25^ (Supplementary Fig. 6c) and most were implicated in brain function by gene ontology analysis (Supplementary Fig. 6d). In addition to this group of genes, a gene set that has been implicated in a variety of neurological disorders^47^ also showed behavior compatible with repression by MeCP2, as the magnitude of mis-regulation in KO and OE hypothalamus correlated reciprocally with MeCP2 occupancy (Fig. 6k-l).

## Discussion

### DNA methylation is the dominant determinant of MeCP2 binding in the brain

This study explores the determinants of MeCP2 binding to DNA *in vitro* and *in vivo* and their relationship to gene expression. It has been shown previously that DNA methylation is the primary determinant of binding in cultured cells by genetic ablation of the DNA methyltransferase Dnmt1^15,27^. In mature neurons, however, where MeCP2 is extremely abundant^16^, MeCP2 binding also tracks DNA methylation^16^, but may depend on other features of chromatin. To rigorously test whether DNA methylation is a pre-requisite for MeCP2 binding in brain, we depleted Dnmt1 in the young brain and tested the effect on MeCP2 binding by ChIP. At this early postnatal stage, mCAC is low and mCG is therefore the predominant modified sequence. The results show clearly that a ∼40% reduction in DNA methylation causes a commensurate loss of chromatin-bound MeCP2 on randomly selected single copy and repetitive genomic sequences and therefore sustains the view that this modification is quantitatively the major determinant of binding.

While this paper was under review a study of MeCP2 binding in olfactory bulb of mouse reported contrasting conclusions^48^. The authors found that MeCP2 is enriched at non-methylated CGIs and that DNA methylation was apparently a minor determinant of binding. This conclusion conflicts with several previously published MeCP2 ChIP experiments from different laboratories, which found a dramatic drop in MeCP2 binding at CGIs in cultured cells^15^, whole mouse brain^16^, hypothalamus^26^, cortex and cerebellum^25^ coupled with DNA methylation-dependent occupancy of the genome. A potential explanation for the discrepant results in olfactory bulb is that the rules governing MeCP2 binding in this region of the brain differ from those operating in the remainder of the nervous system. Alternatively, there may be a technical issue regarding antibody specificity that has led to inconsistent findings. It is notable that the earlier reports showing DNA methylation dependence were achieved using a diversity of validated antibodies.

### DNA methylation-dependent recognition sequences of MeCP2

We comprehensively analyzed the modified DNA sequences that determine MeCP2 binding. To insure the reliability of our conclusions we used three experimental approaches: *in vitro* EMSA; *in vivo* ChIP in transfected cultured cells; and *in vivo* ChIP-seq of mouse brain. Three methylated DNA motifs consistently recruited MeCP2: mCG, mCAC and hmCAC. Interestingly, mCAC is the predominant methylated non-CG sequence in brain, comprising 15 – 30% of all methylated cytosine in sorted mouse neurons, probably due to the action of the *de novo* DNA methyltransferase Dnmt3a^2,4^. The hydroxymethylated sequence hmCAC, on the other hand, is reportedly extremely rare perhaps due to the preference of Tet enzymes for mCG as a substrate^3^. Given the inability of MeCP2 to bind hmCG and the apparent extreme rarity of hmCAC, it seems unlikely that hmC is a major contributor to the biological role of MeCP2.

Our modeling provides a structural explanation for the sequence specificity of the interaction between MeCP2 and DNA. We previously observed that the replacement of a mC at a methylated CG di-nucleotide with T, forming a T:G mispair, had a negligible effect upon the binding affinity of MeCP2^36^, indicating that hydrogen bonding with the mC/T is not essential and that the interaction is flexible enough to accommodate the T:G wobble geometry. Here we observed that in duplex DNA pyrimidine-methyl groups can be provided by either the mC or T, as demonstrated by the finding that the replacement of T with U, which lacks the T methyl group, results in loss of MeCP2 binding. Remarkably the observed specificity for mC in either mCG or mCAC sequence contexts can be rationalized with minimal changes to the conformation of the MBD that was established by X-ray crystallography^40^. Only the configuration of the side-chain of amino acid arginine 133 needs to be adjusted to account for both permitted and non-permitted interactions. The model has potentially important biological consequences, as it suggests that binding to mCAC or mCG is structurally very similar, making it likely that MeCP2 binding to any of these sequences will lead to the same outcome down-stream. In agreement with this scenario, experiments comparing reporters methylated at mCA or mCG suggest that both modified sequences lead to transcriptional repression^2^. Missense mutations that cause Rett syndrome predominantly affect the MBD or the NID, suggesting that the primary role of MeCP2 is to bridge DNA sequences with the NCoR/SMRT corepressor complexes^13^. We propose that mCG and mCAC, function identically to facilitate this bridging process.

### MeCP2 binding profiles and gene expression

The high frequency of mCG and mCAC binding sites for MeCP2 throughout the neuronal genome (one per ∼100 bp) poses problems for conventional ChIP-seq analysis. In most studies binding sites of, for example, transcription factors are widely spaced compared with the size of chromatin-derived DNA fragments generated by ChIP protocols (200-500 bp). Therefore, only a minority of DNA fragments is expected to be immunoprecipitated and discrete peaks are recovered. In the case of MeCP2, most DNA fragments of ∼200-500 bp contain binding sites (CGIs being a conspicuous exception) leading to a relatively uniform recovery of genomic DNA that resembles Input. The difference between Input and ChIP signal, which is the measure of MeCP2 density, provides an undulating continuum in which peaks are broad.

A striking feature of the *Mecp2* KO hypothalamus is the reduced level of total RNA, matching reports in cultured mouse and human neurons^24,46^. The mechanism responsible is unknown, but one possibility is that MeCP2 is a direct global activator of transcription^23,24^. Arguing against this possibility, we found that most KO down-regulated genes lie in domains of low MeCP2 occupancy. Also, it might be expected that two-fold over-expression of an activator would lead to increased levels of RNA compared to WT, but this is not observed (Supplementary Fig. 5a). An alternative explanation is that reduced RNA reflects reduced cell size, perhaps as a secondary consequence of sub-optimal neuronal gene expression. In this case the change in total RNA and the relative mis-regulation of genes within the RNA population may be separate phenomena. Given the close relationship between gene mis-regulation, MeCP2 binding site density and MeCP2 occupancy, we favor the view that the effects of MeCP2 concentration on the balance of neuronal gene expression are primary, whereas the downward shift in total RNA is a secondary effect. To test this rigorously it will be important to track down the origins of the total RNA deficiency.

By re-analyzing ChIP-seq and RNA-seq datasets from hypothalamus of WT mice, *Mecp2*-null mice and mice over-expressing MeCP2 we were unable to confirm reports that MeCP2 is more highly bound to the transcription units of mis-expressed genes regardless of up- or down-regulation^26^. Instead we found that MeCP2 is bound at higher levels within and surrounding genes that are up-regulated when MeCP2 is missing, or down-regulated by MeCP2 over-expression. These findings fit well with the evidence that MeCP2 may link methylated DNA with the NCoR/SMRT corepressor^13^ and they endorse the conclusions of Gabel and colleagues^25^ who found repression of long genes by MeCP2. Over-expression would be expected to increase repression of the most binding site-rich genes whereas MeCP2 depletion would preferentially relieve their inhibition, as is observed. Although we emphasize the correlation with binding site density per unit of DNA length, it is possible that the absolute number of binding sites per gene also contributes to the MeCP2 response. Further work is required to disentangle the roles of these related variables.

A notable feature of the MeCP2 binding site and occupancy profiles is that enriched or depleted regions are larger than genes. Mis-regulated genes belong to large domains that are rich in mCG and mCAC with relatively homogeneous MeCP2 binding levels extending up- and down-stream of the transcription unit. Within one domain, however, not all genes respond in the same way to changes in MeCP2 abundance, perhaps due to additional transcriptional regulatory mechanisms. The tri-nucleotide mCAC, despite its lower abundance compared with relatively uniformly distributed mCG, correlates strongly with MeCP2 binding and transcriptional mis-regulation in response to altered levels of MeCP2. The accumulation of mCAC during development of the brain adds numerically to the number of MeCP2 binding targets in the genome, but it also changes the distribution of binding sites. Our analysis agrees with previous suggestions that mCA is biologically important for MeCP2 function in relation to transcription^25^.

Superficially, mCG correlates less well than mCAC with changes in gene expression. This effect may be exaggerated by the difference in their densities, however. Mis-regulated genes have on average 0.5 extra mCAC per 1000 bp when compared with those changing in the opposite direction. As the density of mCG in the genome is higher than mCAC (∼15 per kb), a comparable incremental difference in mCG density would represent <4% of the total and would not therefore be reliably measurable. These considerations may mean that the small differences in gene expression in response to changing amounts of MeCP2 are accompanied by equivalent subtle differences in the densities of both mCG and mCAC. Which bound MeCP2 could mediate repression of this kind? It is possible that only MeCP2 occupancy within a gene is relevant to regulation, the flanking methylation playing no role. Alternatively, the regulatory influence of MeCP2 may operate over large domains encompassing many genes. Future research will attempt to distinguish these possibilities.

### Subtle effects on transcription of many genes

While there is no direct evidence that aberrant gene expression is the proximal cause of Rett syndrome or MeCP2 over-expression syndrome, it is noteworthy that thousands of genes, including many implicated in human neuronal disorders, are sensitive to altered levels of MeCP2. Mild mis-regulation on this scale may destabilize neuronal function^25^. It is worth recalling that Rett neurons, though sub-optimal, are viable for many decades. In this sense the biological defect can be seen as mild, despite the profound effects on higher functions of the brain. The challenge now is to determine how brain function might be affected by a multitude of small discrepancies in gene expression. Overall, the results presented here sustain a coherent view of MeCP2 function: namely that MeCP2 binding at mCG and mCAC sites determines the magnitude of a repressive effect on transcription that is exacerbated by MeCP2 excess and relieved by MeCP2-deficiency. With the benefit of a comprehensive list of MeCP2 target sequences at the molecular level, the predictions of this model can be experimentally tested, clarifying further the role of MeCP2 in regulating transcription in the brain.

## Online Methods

### Animal care and transgenic mouse lines

All mouse studies were approved by the Austrian Animal Care Committee and were licensed under the UK Animals (Scientific Procedures) Act 1986 and conducted in accordance with guidelines for use and care of laboratory animals. To delete *Dnmt1* in the nervous system, mice with a floxed *Dnmt1* allele^49^ were mated with Nestin-Cre mice^50^. The *Dnmt1* Nestin-Cre strain was bred in a mixed genetic background C57BL/6J x 129SV. Male *Mecp2*^*STOP/y*^ and corresponding WT littermates^51^ and male *Mecp2* -/y and WT littermates^31^ were used as Western blotting, Real Time PCR and ChIP controls. C57Bl6 male WT 10 week old mice were used for FACS sorting experiments and consecutive WGBS and TAB-seq.

### Normalization of mRNA

Hypothalamus was isolated from 6 week old male WT and *Mecp2* -/y mice in 5 replicates. RNA and DNA were co-isolated with the AllPrep DNA/RNA Mini kit *(Qiagen)* according to manufacturer’s instructions with some modifications. In short, tissue was homogenized in 1ml RLT buffer (spiked with 3 × 10^6^ *Drosophila* S2 cells/10ml RLT buffer) and centrifuged in a *Qiashredder* column *(Qiagen)* for 2 minutes at full speed. The eluted RNA was next subjected to treatment with the *DNA-free* DNA removal kit *(Ambion)* and reverse transcribed with the iScript cDNA synthesis kit *(BioRad).* Real Time quantitative PCR was performed on cDNA and DNA with *Drosophila* and mouse specific mRNA and genomic DNA primers. For analysis, mouse mRNA was normalized to *Drosophila* RNA and analogous mouse DNA was normalized to *Drosophila* DNA. In the final step, corrected mRNA levels were normalized to corrected DNA values. Primer sequences can be found in Supplementary Table 3.

### Mouse brain nuclei isolation and FACS sorting

Brain nuclei isolation and consecutive FACS sorting according to NeuN expression was performed as described previously^16^. 10 week old WT Bl6 male mice were used and 4 brains were pooled for each replicate.

### Nuclear protein extracts and Western blotting

Whole brain protein extracts were prepared as previously described with modifications^13^. After homogenization in NE10 buffer and addition of 250 units Benzonase *(Sigma),* equal amounts of 2x SDS Loading Buffer were added and extracts boiled for 3 minutes. Western blotting was performed according to standard procedures. Antibodies used for Western: DNMT1 1:1000 (gift from Dr. Amir Eden); MeCP2 M6818 (C-terminal) 1:1000 *(Sigma);* MeCP2 M7443 (N-terminal) 1:1000 *(Sigma);* β-Actin A5316 *(Sigma)* 1:5000; Gapdh D16H11 *(Cell Signaling)* 1:5000.

### Verification of modified oligonucleotides

Dot blots of modified oligonucleotides and control DNA (Methylated standard kit, *Active Motif)* were generated with Bio-Dot^®^ Microfiltration Apparatus *(BioRad)* using manufacturer’s recommendations. Oligonucleotides and control DNA were denaturated by the addition of [0.4M] NaOH, [10mM] EDTA in a total volume of 50μl and boiled for 10 minutes. DNA was neutralized by addition of an equal volume of ice-cold 2M ammonium acetate. Control DNA and oligonucleotides were spotted in duplicate serial dilutions (Control DNA: 50ng, oligonucleotides: 10μM starting concentration). Nitrocellulose membrane was UV auto-crosslinked and then blocked for 30 minutes in 5% non-fat dried milk powder, 0.05% *Tween* 20/ 1x TBS. Primary antibodies were incubated for 45 minutes at room temperature (5hmC: 1:10.000 *Active Motif;*5mC: 1:500 *Eurogentech).* Secondary *LI-COR* antibodies were incubated in the dark for 30 minutes (donkey anti mouse IRDye 800Cw; donkey anti rabbit IRDye 680; *LI-COR).* Membranes were scanned with a *LI-COR* Odyssey instrument.

### Protein expression, purification and EMSA

Protein was prepared as described^39^. When examining MeCP2 [1-205] specificity, DNA sequence was varied at the tri-nucleotide indicated in bold (See Supplementary Table 1). The primary cytosine of this tri-nucleotide was either non-methylated, methylated, or hydroxymethylated. All oligonucleotides were annealed to their complement, ^32^P-labelled and electrophoretic mobility shift assays performed on ice for 30 min using conditions described previously^43^. In competition assays to assess tri-nucleotide-binding preferences of MeCP2 [1-205] a parent 58 bp *Bdnf-probe,* containing the centrally methylated sequence mCGG, was ^32^P-labelled and co-incubated with a 2000-fold excess of cold-competitor DNA bearing one of the sequences described in Fig. 2b-c. Bound complexes were resolved as described above and levels of competition visualized by Phosphorimager analysis and ImageJ quantification. These experiments were performed in triplicate.

### Preparation of total nucleic acid for estimation of RNA versus DNA quantity

Dissected hypothalamus tissue was homogenized in lysis buffer (10mM Tris HCl [pH 7.4], 0.5% SDS, 100mM EDTA, 300μg/ml proteinase K) and incubated at 50°C for 2 hours. Total nucleic acid was recovered from the completely lysed sample by ethanol precipitation in 2 volumes of 100% ethanol at room temperature (for 30 minutes), and pelleting by centrifugation. The pellet was washed once in 2 volumes of 70% ethanol, and the nucleic acid pellet was resuspended in hydrolysis buffer containing 1x DNase I buffer *(NEB),* 1mM zinc sulphate, DNase I *(NEB)* and Nuclease P1 *(Sigma).* After 4 hours the sample was mixed thoroughly and digested for a further 8 hours. After 12 hours at 37°C, the sample was heated to 92°C for 3 minutes and cooled on ice. Two volumes of 30mM sodium acetate, 1mM zinc sulphate [pH 5.2] were added plus additional Nuclease P1 and the nucleic acids were further digested to deoxyribonucleotide and ribonucleotide 5’ monophosphates for a further 24 hours at 37°C. The samples were then subjected to HPLC as set out below.

### HPLC estimation of 5mC

Measurement of genomic cytosine methylation in mouse brains was as described^52^. After DNA extraction residual RNA was removed by enzymatic hydrolysis (6-hour incubation with RNase A and RNase T1) followed by DNA precipitation in 3 volumes of ethanol. This procedure was repeated once. Purified DNA was digested with DNase I *(NEB)* for 12 hours in the recommended buffer after which the sample was heated (92°C for 3 minutes) to denature any remaining double stranded DNA. After cooling on ice 2 volumes of 30mM sodium acetate, 1mM zinc sulphate [pH 5.2] was added and the DNA was further digested to nucleotide 5’ monophosphates with Nuclease P1 *(Sigma)* for 12 hours. The quantification of 5-methylcytosine in genomic DNA was by isocratic high performance reverse phase liquid chromatography as previously^52^, with the following alterations. A Dionex UM 3000 HPLC system was used complete with a column chiller, C18 column (250mm x 4.6mm 5 um APEX ODS, #4M25310, *Grace Discovery Sciences),* and column guard *(Phomenex,* #AJ0-7596). The mobile phase was 50mM (monobasic) ammomium phosphate [pH4.1]. The column was chilled to 8°C to improve peak separation. Deoxyribonucleotides (dNMPs) were detected over a 70 minute run time using a Dionex 3000 multiple wavelength detector.

### HPLC nucleotide quantifications

UV absorbance was recorded at 276 nm (dCMP, elution time 9.4 minutes), 282 nm (5mdCMP, elution time 17 minutes), 268 nm (dTMP, elution time 21.9 minutes), 260nm AMP and dAMP (elution times 27 minutes and 62.47 minutes) and 254 nm (GMP and dGMP, elution times 11.1 minutes and 29.7 minutes). Extinction coefficients used in nucleotide quantifications were dCMP, 8.86 × 10^3^; 5mdCMP 9.0 × 10^3^; dTMP, dGMP/GMP 12.16 × 10^3^; dAMP/AMP 15.04 × 10^3^. Quantifications were calculated from the area under each peak estimated using Chromeleon software using the respective extinction coefficients.

### Transfection ChIP assay

HPLC purified oligonucleotides and corresponding antisense oligonucleotides were purchased from *biomers.net.* Some oligonucleotides containing 5hmC were synthesized and characterized as described previously^53^. All oligonucleotide sequences used are listed in Supplementary Table 1. Equal amounts of sense and antisense oligonucleotide stocks (100μM) were mixed with 10x Ligation Buffer *(NEB)* in 50μl volumes. Oligonucleotide mix was boiled in a water bath for 8 minutes and cooled to room temperature. Annealed oligonucleotides were cleaned up with MSB Spin PCRapace cleanup kit *(Invitek)* and diluted to 10μM stocks for transfections. HEK293FT cells (1.5 × 10^6^) were transfected overnight with 0.5μg of full length MeCP2 tagged at the N-terminus with EGFP using Lipofectamine 2000 *(Lifetechnologies)* according to manufacturer’s instructions. After assessment of transfection efficiency (described in ^39^), the medium was changed and replaced with annealed unmodified, methylated or hydroxymethylated oligonucleotides [100nM final concentration] using TransIT Oligofect reagent *(Mirus)* for 4 hours. Cells were washed with PBS and harvested by scraping. Cells were then crosslinked with formaldehyde to 1% final concentration for 5 minutes at room temperature and quenched by the addition of glycine to a final concentration of 0.125M for 5 minutes followed by another two washes in PBS. Cell pellets were flash frozen in liquid nitrogen and stored at -80°C or directly used for chromatin isolation and consecutive ChIP with 4μg MeCP2 M6818 antibody *(Sigma).* Primer sequences can be found in Supplementary Table 2.

### Mouse brain chromatin preparation

Frozen or fresh mouse brains were homogenized in PBS supplemented with Protease Inhibitor Cocktails *(Roche)* and crosslinked with formaldehyde to 1% final concentration for 10 minutes at room temperature. After quenching the crosslink with 0.125M glycine, brains were washed in ice cold PBS twice and soluble chromatin preparation was performed.

### Chromatin preparation and ChIP

Soluble chromatin preparation and ChIP assays were carried out as described previously^54^ with some modification. In short, chromatin was sonicated using a Twin Bioruptor *(Diagenode)* 30sec on/off for 15 cycles at 4°C. Equal amounts of chromatin were used for IP with 4μg MeCP2 M6818 antibody *(Sigma)* and incubated overnight. Protein-antibody complexes were bound to magnetic protein G beads *(Lifetechnologies)* for 4-5 hours and washed with standard IP wash buffers for 10 minutes at 4°C. The crosslink was reversed by addition of 0.05 volume of 4M NaCl overnight at 65°C. After proteinase K digestion, DNA was recovered by phenol-chloroform-isoamylalcohol extraction and dissolved in 200μl H_2_O. Real Time PCR of diluted ChIP DNA and corresponding Input DNA was performed on LightCycler (*Roche*). All primer sequences used for ChIP are listed in Supplementary Table 2.

### Structural modeling

Modeling was based on the X-ray structure of MeCP2 (PDB code 3C2I) using the program COOT^55^. Atomic coordinates for DNA bases were generated using the ‘mutate’ option. To optimize hydrogen-bonded and van der Waals contacts between protein and different base pair sequences, the conformation of the side chain of R133 was adjusted manually (all other atoms in the structure were left unchanged). The potential role of water molecules in the recognition of different base-pair sequences by MeCP2 was examined by placing a water molecule in the highly conserved and most probable sites of hydration in the major groove of B-DNA as described^56^. All figures were prepared using the graphics program PyMol (*DeLano Scientific, San Carlos, CA*).

### Locus-specific bisulfite sequencing

Bisulfite treatment of genomic DNA of P11 WT and *Dnmt1* KO whole brain triplicates and DNA sequencing was carried out as previously described^57^ with some modifications. Genomic DNA was isolated with the DNeasy Blood & Tissue kit (*Qiagen*) and bisulfite treated with the EpiTect bisulfite kit (*Qiagen*). Bisulfite Primers were designed with MethPrimer software^58^. Sequences were analyzed with the BISMA online tool^59^.

### Reduced representation bisulfite sequencing

Genomic DNA from whole brain of 5 day old *Dnmt1* WT and KO littermate replicates was isolated using the DNeasy Blood & Tissue Kit (*Qiagen*). For RRBS^34^, the “Reduced Representation Bisulfite Sequencing for Methylation Analysis” protocol (www.support.illumina.com) was followed and TruSeq^®^ v2 DNA sample preparation kit *(Illumina)* was used. Cleaned up libraries were validated on a Bioanalyzer High Sensitivity DNA Chip *(Agilent)* and 100 bp paired-end sequencing was performed on Illumina HiSeq 2000 platform (Wellcome Trust Sanger Institute, Hinxton, UK).

### Whole genome bisulfite and TAB sequencing

Genomic DNA of three replicates of 10 week old male Bl/6 WT dissected hypothalamus samples was prepared with the DNeasy Blood & Tissue Kit *(Qiagen)* and 0.5% unmethylated λ DNA *(Promega)* was spiked in. Equal amounts of genomic DNA were bisulfite converted with the EZ DNA Methylation Gold Kit *(Zymo Research)* and libraries were prepared with the TruSeq DNA methylation kit *(Illumina)* according to manufacturer’s instructions. WGBS and TAB sequencing from NeuN positive sorted neuronal nuclei was as described previously^60^. For TAB treatment, half the DNA was glycosylated, TET oxidized and spiked with control DNA. The other half was left untreated and spiked with unmethylated *λ* DNA *(Promega).* NGS libraries were prepared with TruSeq DNA Sample Preparation Kit *(Illumina)* according to manufacturer’s instructions. After size selection, all libraries were bisulfite treated with EpiTect Bisulfite Kit (*Qiagen*) and amplified with Pfu Turbo Cx Polymerase *(Stratagene)* for 7 PCR cycles. Cleaned-up libraries were validated on a Bioanalyzer High Sensitivity DNA Chip *(Agilent)* and 100 bp paired-end sequencing performed on an Illumina HiSeq 2000 platform (Wellcome Trust Sanger Institute, Hinxton, UK).

## Bioinformatics and statistical analyses

### Genome build and annotations

All data were aligned to the mouse NCBI 37 (mm9) assembly. Gene annotations were obtained from version 67 of the Ensembl database.

### ChIP-seq

MeCP2 Chip and Input datasets were downloaded from the Gene Expression Omnibus (GEO) under the accession numbers GSE66868^26^ and GSE60062^25^. Raw sequencing reads were first trimmed and filtered using Trimmomatic v0.32^61^ then aligned with bwa using the samse algorithm. Alignments were then filtered to remove reads classed as duplicates, non-unique or those that fell in the blacklisted regions outlined by the Encode project^62^. Alignments were converted to bigWig files using the deepTools package for genome wide visualization and analysis. Read counts were normalized to RPKM to account for differences in library size. The bigwigCompare tool was used to calculate a log2 ratio of MeCP2 ChIP/Input signal across the genome.

### Whole genome bisulfite sequencing of mouse hypothalamus

Quality trimming, filtering and adapter removal were performed by trimmomatic v0.32 prior to alignment. The software package Bismark v0.15 was used to map reads to the mm9 genome, remove duplicate alignments and extract methylation calls^63^.

### BS-seq and TAB-seq

Processed bisulfite and TAB-seq datasets containing aligned methylation calls were obtained from the GEO records GSM1173786_allC. MethylC-Seq_mm_fc_male_7wk_neun_pos and GSM1173795_allC.TAB-Seq_mm_fc_6wk respectively. Percentage (%) DNA methylation at a given site *i* corresponds to the ratio of mC basecalls for that site to the count of all reads mapping to that site (*m^i^* = *mC/C*). Context-specific % mean methylation for a given region (i.e. bin or gene) was defined as 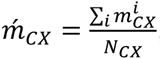, where *N_cx_* is the number of C’s within the region, occurring in context *CX.* In addition, context-specific methylation density was defined as 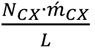, where L is the length of the region.

### RNA-seq

Raw reads from mouse hypothalamus were downloaded from the Gene Expression Omnibus (GEO) under the accession no. GSE66870^26^. Trimmomatic v0.32 was used to remove adaptor contamination and to trim low quality reads. Reads were mapped to the genome using STAR v 2.4.2a^64^. Alignments were then filtered to remove non-unique and blacklisted reads. HTseq-count v0.6.0 was used to quantify read counts over gene exons in the union mode.

### Correlation between MeCP2 ChIP-Seq and Input read counts (Fig. 5e-f)

For the MeCP2 ChIP and Input sample of the Chen dataset^26^ we computed the number of reads that mapped to 10 kb windows covering the entire genome. For this we have shifted the reads by 134 bp which correspondents to the estimated fragment length determined by MACS. A Kernel density estimate was applied. We have applied linear regression (MeCP2 ∼ Input) and analysed the residuals *r_i_* for each bin. Windows with residuals exceeding, a threshold are considered enriched or depleted respectively. The threshold was chosen to be 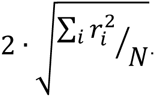

### Correlation between log2(MeCP2/Input) and DNA methylation in genomic windows (Fig. 5i-j and Supplementary Figs. 4j-r)

We calculated read coverage values and context-dependent methylation densities for 1 kb windows across the genome. Regions were ordered by their log2(MeCP2 Chip/Input) signal and mean methylation densities were calculated for groups of 1000 regions. Datasets used are ^26^ and hypothalamus WGBS (this study).

### MeCP2 summit analysis (Fig. 5g-h and Supplementary Figs. 4b-i)

Summits of MeCP2 ChIP enrichment over Input were defined using MACS and the method described in ^25^ for both MeCP2 datasets (^25,26^). We used the Bioconductor package *seqplots v1.4.0* to plot methylation density across the summits for various cytosine contexts. Mean methylation densities were calculated for 20 bp windows across the summits extended by 4 kb on either side. The ChIP-seq dataset from cortex was combined with bisulfite analysis from ^3^ (Fig. 5g and Supplementary Figs. 4i, l-m) and the hypothalamus ChIP-seq dataset ^26^ was correlated with bisulfite data from hypothalamus WGBS (this study) (Fig. 5h and Supplementary Figs. 4b-h and n-r) or bisulfite data from ^3^ (Supplementary Figs. 4j-k).

### Rolling mean plots for genic regions (Fig. 5k)

Genes were sorted according to their MeCP2/Input enrichment and rolling means of methylation density were applied over subsets of 400 genes with a step of 80 genes. Datasets used are ^26^ and hypothalamus WGBS (this study).

#### MSR (Fig. 5l-m and Supplementary Figs. 4s-u)

We used the MSR tool to find domains of MeCP2 enrichment and depletion relative to the Input sample^42^. As background we used a mappability map for the m9 genome (parameters: L=45, P-value threshold: 1e-6). For each significantly enriched or depleted segment we determined the methylation densities and averaged over segments with same scale and enrichment score. Datasets used are ^26^ and hypothalamus WGBS (this study).

#### Differential expression analysis (Fig. 6)

We used DeSeq2 v1.8.1^65^ to determine mis-regulated genes in KO and OE. To account for the observed 15% reduction of total mRNA in the KO samples, we first used the DESeq function estimateSizeFactors to normalize the data sets and subsequently multiplied the obtained size factors for the KO samples by 1.15. An adjusted p-value threshold of <0.05 was used to determine up- or down-regulated genes. For an exemplary group of genes for which little change is observed between conditions we used p_adj >0.5 with a log2FoldChange <0.01. We further filtered out protein-coding genes with constitutively low expression (TPM <5 in all samples) and genes with the lowest Input coverage. We retained 12,510 protein-coding genes, of which 403 showed higher, and 8363 lower expression in KO vs WT, respectively. In addition, we found 1372 to have higher and 2719 to have lower expression in OE relative to WT. Dataset used is ^26^

#### Aggregated signal plots across gene features (Fig. 6a-f and Supplementary Fig. 4a)

We used *seqplots* to generate aggregate plots at transcription start sites of protein coding genes and annotated CGIs. Mean values for log2(MeCP2 ChIP/Input) and methylation densities were calculated over 1 kb windows for each set of genes and 100 bp windows for CGIs (Supplementary Fig. 4a). Datasets used are ^26^ and hypothalamus WGBS (this study).

## Author Contributions

SL, JCC and GSch performed or directed experiments and analyses in the study. SL designed and performed *Dnmt1* KO mouse experiments and BHR did HPLC. JCC and MDW designed and performed EMSA analyses and structural modeling. SL and LS designed and performed *in vivo* transfection assays. SL, JCC and DD performed bisulfite sequencing and fragment cloning. SL, CH and MY designed and performed WGBS and TAB sequencing. GSch, SW and GSi performed bioinformatics and statistical analyses. JS and CS provided mouse reagents. LS provided 5hmC-containing oligonucleotides and contributed ideas. AB, SL, JCC and GSch wrote the manuscript, SL prepared the figures and all co-authors passed comment. AB advised on all aspects of the study.

## Acknowledgements

We are grateful to Kashyap Chhatbar for bioinformatics assistance and to Michael Greenberg, Gail Mandel and their group members for valuable Input. We also thank Justyna Cholewa-Waclaw, Rebekah Tillotson, Ruth Shah and Martha Koerner for useful discussions and Astrid Hagelkruys, Alan McClure and Martin Waterfall for technical assistance. We are grateful to Amir Eden for the DNMT1 antibody and the Patrick Heun lab for providing *Drosophila* S2 cells and primers. The work was supported by a Consortium Grant from The Rett Syndrome Research Trust and by a Wellcome Trust programme grant (091580), Wellcome Trust Centre Core Grant (092076), European Research Council (MLCS 306999) (GSi) and Austrian Science Fund (P25807) (CS). SL and GSch held EMBO long-term fellowships. GSch was also funded by a Marie Curie Postdoc fellowship and SL is currently a fellow of the Veterinary Medicine University of Vienna Postdoc program. LS was supported in part by grant CA184097 from the National Cancer Institute, National Institutes of Health.

